# A Library of Custom PEG-Lipids reveals a Double-PEG-Lipid with Drastically Enhanced Paclitaxel Solubility and Human Cancer Cell Cytotoxicity when used in Fluid Micellar Nanoparticles

**DOI:** 10.1101/2024.08.01.606138

**Authors:** Aria Ghasemizadeh, Lili Wan, Aiko Hirose, Jacqueline Diep, Kai K. Ewert, Cyrus R. Safinya

**Affiliations:** Materials Department, University of California, Santa Barbara, California 93106, USA; Biomolecular Science and Engineering, University of California, Santa Barbara, California 93106, USA; Molecular, Cellular, and Developmental Biology Department, University of California, Santa Barbara, California 93106, USA; Physics Department, University of California, Santa Barbara, California 93106, USA

## Abstract

Paclitaxel (PTX) is one of the most widely utilized chemotherapeutics globally. However, the extremely poor water solubility of paclitaxel necessitates a mechanism of delivery within blood. Fluid lipid PTX nanocarriers (lipids in the chain-melted state) show promise as PTX delivery vectors, but remain limited by their solubility of PTX within the membrane. To improve pharmacokinetics, membrane surfaces are typically coated with polyethylene glycol (PEG). Recent work has demonstrated the generation of a population of micelles within fluid lipid formulations containing a 2kDa PEG-lipid at a 10 mol% ratio. Driven by the positive curvature of the PEG-lipid (i.e. area of head group > area of tails), micelle-containing formulations were found to exhibit significantly higher uptake in cancer cells, cytotoxicity, and *in vivo* antitumor efficacy compared to formulations containing solely liposomes. Here, we describe the custom synthesis of a library of high-curvature micelle-inducing PEG-lipids and examine the effects of PEG chain length, chain branching (single- or double-PEG-lipid), and cationic charge on PTX solubility and cytotoxicity. We examined PEG-lipids at standard (10 mol%) and high (100-x mol%, where x=PTX mol%) formulation ratios. Remarkably, all formulations containing the synthesized high-curvature PEG-lipids had improved PTX solubility over unPEGylated formulations and commercially available DOPE-5k. The highest PTX solubility was found within the 100–xptx mol% PEG-lipid micellar formulations, with particles made from 2k_2_ (two PEG2k chains) encapsulating 13 mol% PTX for up to 24 h. The pancreatic cancer cell line PC3 exhibited higher sensitivity to formulations containing PEG-lipid at 100–xptx mol%, the most potent of which being formulations made from 2k_2_ (IC50 = 14 nM). The work presented here suggests formulations employing high-curvature PEG-lipids, particularly the double-PEG-lipid 2k_2_, hold great potential as next-generation PTX delivery systems owing to their high PTX solubility, enhanced cell cytotoxicity, and ability for precision targeting by affixation of ligands to the PEG molecules.

## Introduction

Paclitaxel (PTX) is a hydrophobic (LogP = 3.66) small molecule drug first isolated from the bark of the Pacific Yew in 1962.^1, 2^ Within cells, PTX functions as a stabilizer of microtubules, binding β-tubulin with high affinity and resulting in reduced dynamic control of microtubule polymerization.^3^ Initially approved for treatment of ovarian cancer in 1992, today PTX is used as a first-line therapy for numerous cancers including breast, bladder, lung, prostate, esophageal, and Kaposi sarcoma.^4^ PTX has been shown to induce cancer cell death by interfering with mitotic spindle formation, causing mitotic arrest that leads to apoptosis.^5, 6^ Since its initial discovery, the efficacy of PTX as an anticancer drug has been severely hindered by its poor solubility in water (<0.25 µg/mL), necessitating its co-administration with excipients such as Cremophor EL (1:1 v/v polyethoxylated castor oil:dehydrated ethanol) in a formulation referred to as Taxol^®^.^7^ However, Cremophor EL introduces its own complications, with a subset of Taxol patients being documented as suffering nephro- and neurotoxicity in addition to hypersensitivity reactions such as anaphylaxis, hypotension, and angioedema.^8, 9^ These side effects are compounded by the low PTX loading (1 wt% relative to Cremophor EL), requiring high doses and long infusion times.^10^ The clinical demand for improved PTX formulations resulted in development of albumin-bound PTX, referred to as Abraxane^®^.^11^ The exclusion of Cremophor EL and the higher PTX loading capacity of Abraxane enhanced the maximum tolerated dose, and Abraxane was approved by the FDA in 2005 for treatment of breast cancer.^12^ Although Abraxane significantly improves breast cancer response rates, it also exhibits faster blood clearance compared to Taxol and has mixed improvement in side effects.^13, 14^ Limiting the effectiveness of all PTX delivery methods is phase separation of PTX from its carrier and formation of crystalline aggregates.^15^ This is undesirable as it results in reduced proportions of pharmacologically active PTX in addition to potential embolization of capillaries by the PTX aggregates.^16–18^

Lipid-based PTX nanocarriers have been developed in an effort to improve upon existing treatments and achieve high PTX loading, long blood circulation times, enhanced tumor accumulation, and minimal off-target effects.^10, 19^ Particles with high PTX loading ratios are advantageous for clinical applications as therapeutic doses require administration of less lipid material which reduces the risk of carrier-induced side effects and therefore increases the maximum tolerated dose, while also lowering the manufacturing cost of the lipid carrier.^20, 21^ The only lipid-based PTX formulation currently approved for clinical use is Lipusu^®^ (approved in China in 2003), with another formulation, EndoTAG-1^®^, currently undergoing Phase III clinical trials for pancreatic cancer in China.^22, 23^ While the composition of Lipusu has not been disclosed, EndoTAG-1 contains the univalent cationic lipid 2,3-dioleyloxypropyltrimethylammonium chloride (DOTAP), neutral 1,2-dioleoyl-*sn*-glycero-3-phosphatidylcholine (DOPC), and PTX at a 50:47:3 mol ratio.^15^ Both DOTAP and DOPC contain monounsaturated dioleyl tails. Lipids with mono and polyunsaturated tails, i.e. lipids forming fluid state (melted-chain) membranes, stably solubilize higher ratios of PTX compared to lipids with saturated tails (with membranes in the gel state) or rigid molecules such as cholesterol.^24–26^ The cationic lipid DOTAP is well-known to promote extravasation *in vivo* by its electrostatic interactions with the anionic sulfated proteoglycans on the surface of endothelial cells.^27–34^ However, DOTAP and other cationic lipids introduce issues of reduced liposome blood half-life, off-target toxicity, and inflammation.^35, 36^ Additionally, traditional liposome formulations solely utilizing monounsaturated diacyl phospholipids such as EndoTAG-1 can typically only stably incorporate PTX at a maximum of 3– 4 mol%.^37–39^ Therefore, while EndoTAG-1 serves as an important benchmark for liposomal PTX carriers, there remains a demand for improved lipid-based PTX nanocarriers with prolonged blood half-life, efficient accumulation in tumor tissue, deep tumor penetration, and stable solubilization of large PTX payloads.^24, 40^

Lipid micelles hold great potential as PTX carriers owing to their size (ca. 10–40 nm) that is an order of magnitude smaller than liposomes (ca. 100–500 nm).^41–44^ Small particle sizes better evade recognition by tissue-resident macrophages, resulting in longer blood circulation time, in addition to improved extravasation from endothelial walls into tumor tissues and deeper tumor accumulation.^45–47^ Yet despite these advantages, to date, no micellar PTX carriers have been tested in clinical trials. Therefore, there is significant merit in developing such carriers.^22, 24, 48^

Many prior micelle preparation methods utilize lysolipids, but are unsuitable for drug delivery purposes owing to their high critical micelle concentrations in the millimolar range that can result in micelle dissolution upon dilution in blood.^40, 49^ The hydrophilic polymer polyethylene glycol (PEG) dramatically prolongs blood circulation time when conjugated to the surface of nanoparticles. When attached to the head group of a diacyllipid, PEG promotes formation of micelle structures because the large volume occupied by the polymer chain results in a positive spontaneous curvature.^50^ Studies have been performed on PEGylated PTX-loaded micelles composed of 1,2-distearoyl-*sn*-glycero-3-phosphoethanolamine (DSPE)-mPEG (with saturated tails) and 1,2-dioleoyl-*sn*-glycero-3-phosphoethanolamine (DOPE)-mPEG (with unsaturated tails) of varying chain lengths.^40, 51–53^ These formulations generated micelles with small particle sizes (<40 nm) and accordingly exhibited robust extravasation and tumor accumulation in mice, but suffered from low PTX incorporation. More recently, a population of micelles was found to coexist with liposomes in cationic PTX formulations containing only 10 mol% of a 2kDa PEG-lipid.^15^ Critically, this PEGylated formulation containing a mixture of liposomes and micelles enhanced cancer cellular uptake and cytotoxicity when compared to the purely liposomal unPEGylated EndoTAG-1 formulation. This effect was also seen *in vivo* with formulations containing 10 mol% DOPE-mPEG5k, demonstrating higher blood half-life, tumor penetration, and apoptotic activity compared to the unPEGylated EndoTAG-1 formulation as well as formulations with only 2 or 5 mol% PEG-lipid.

Modulation of lipid shape can drive selective formation of lipid structures, such as liposomes or micelles.^54^ The spontaneous assembly of lipid structures are dictated by the curvature of the lipid (C_0_), which refers to the ratio of the area of the lipid head group to the cross-sectional area of the tail(s) (Figure 1).^26^ Lipids such as 1,2-oleoyl-*sn*-glycero-3- phosphoethanolamine (DOPE) possess negative curvature (C_0_ < 0) and form inverse micelle structures. Lipids with curvatures close to zero (C_0_ ≈ 0), such as DOPC and DOTAP, form bilayers. Lipids with positive curvatures (C_0_ > 0), including PEG-lipids, form micelles. Spontaneous formation of micelles by lipids with positive C_0_ occurs as a result of the lower bending cost associated with assembly into a micelle compared to a liposome (where local membrane curvature is close to zero).^55–58^

**Figure 1.**
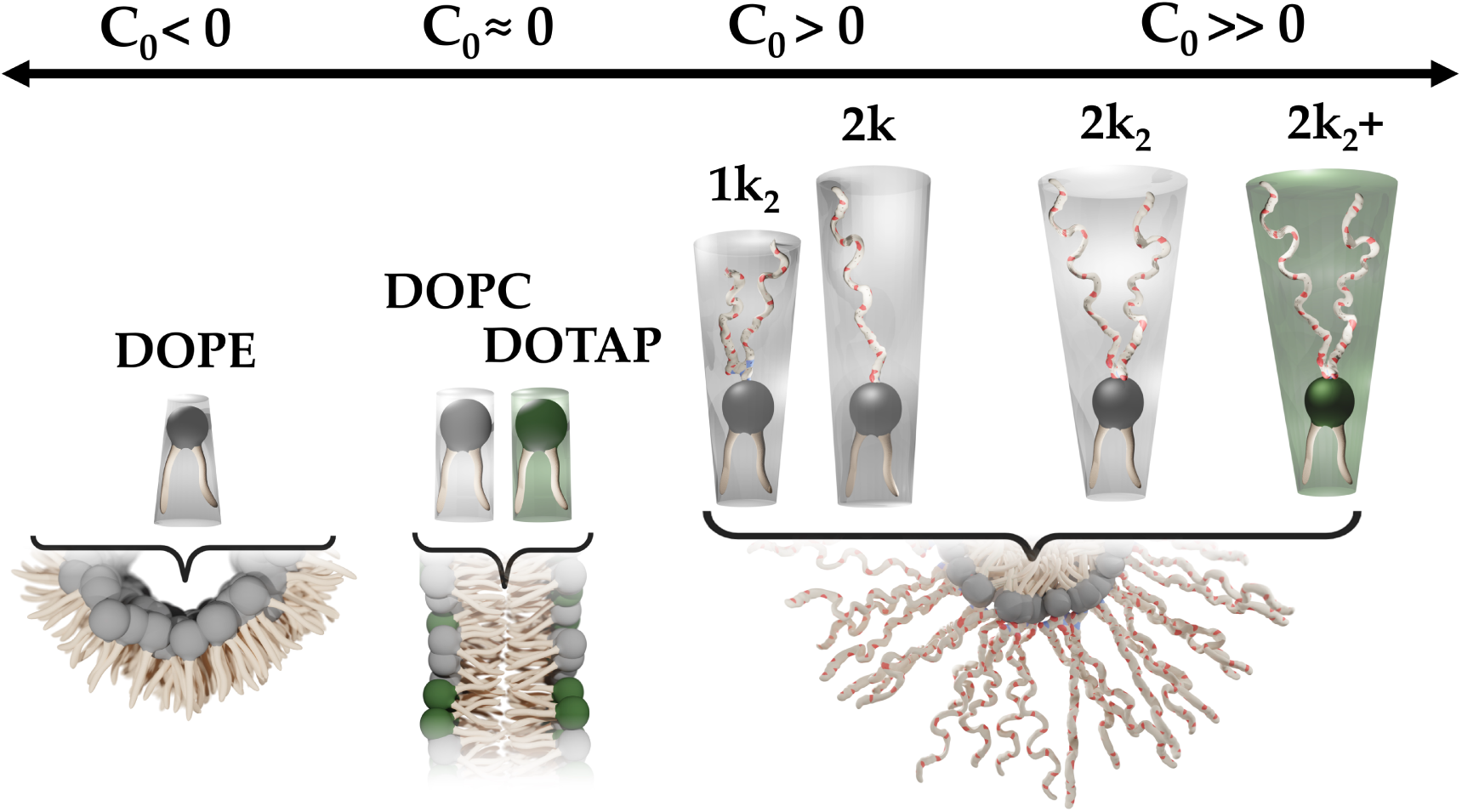
The relationship of lipid curvature and spontaneous assembly of lipid structures. Lipid curvature reflects the ratio between the area of the lipid head group and the tail(s). Lipids with negative curvature (C_0_ < 0) such as DOPE have head group sizes smaller than the tail(s) and form inverted micelles. Lipids with zero curvature (C_0_ **≈** 0) such as DOPC and DOTAP have similarly sized head group and tail(s) and form bilayers. Lipids with positive curvature (C_0_ > 0) such as those synthesized in this study (1k_2_, 2k, 2k_2_, 2k_2_+) have head group sizes larger than the tail(s) and form micelles.

In this study, we sought to synthesize a library of custom high-curvature PEG-lipids for fluid-membrane PTX carriers. These PEG-lipids differ in PEG chain length, PEG chain branching (single or double chain), and lipid charge (neutral or cationic). We elucidated the effects of these parameters on PTX solubility and cytotoxicity in a prostate adenocarcinoma (PC3) cell line. Starting with basic building blocks, we separately synthesized a set of PEG and lipid intermediates, which we then conjugated in an efficient and combinatorial fashion to afford 6 high-curvature PEG-lipids for characterization. For our experiments, we examined the PEG-lipids within two formulation classes (Figure 2): one containing PEG-lipid at 10 mol% (PEG lipid:DOTAP:DOPC:PTX=10:50:40–xptx:xptx mol ratio), which was expected to generate a population of primarily liposomes, and one containing PEG-lipid at 100–xptx mol%, which was expected to generate primarily micelles. We used DIC microscopy to create kinetic phase diagrams (KPD) examining the PTX drug loading solubility and membrane stability over time in formulations containing each PEG-lipid. Remarkably, although PEGylation of liposomes has been reported to reduce PTX loading,^59, 60^ we found higher PTX solubility for the 6 synthesized PEG-lipids in both formulation classes compared to the unPEGylated EndoTAG-1 formulation. Additionally, we discovered higher PTX solubility for the 6 PEG-lipids in both formulation classes compared to commercially available DOPE-mPEG5k (DOPE-5k). Among the two formulation classes, we found the micellar 100–xptx mol% PEG-lipid formulations to solubilize higher ratios of PTX across all 6 PEG-lipids. Increasing PEG-lipid chain length resulted in improved PTX solubility, branching had mixed effects on solubility, and introducing a cationic charge to the PEG-lipids reduced PTX solubility. PEG-lipid formulations that showed high PTX solubility also showed high cytotoxicity. The double-PEG-lipid 2k_2_ was found to be the most potent, with an IC50 = 14 nM, compared to 24 nM for the commercially available DOPE-5k.

**Figure 2.**
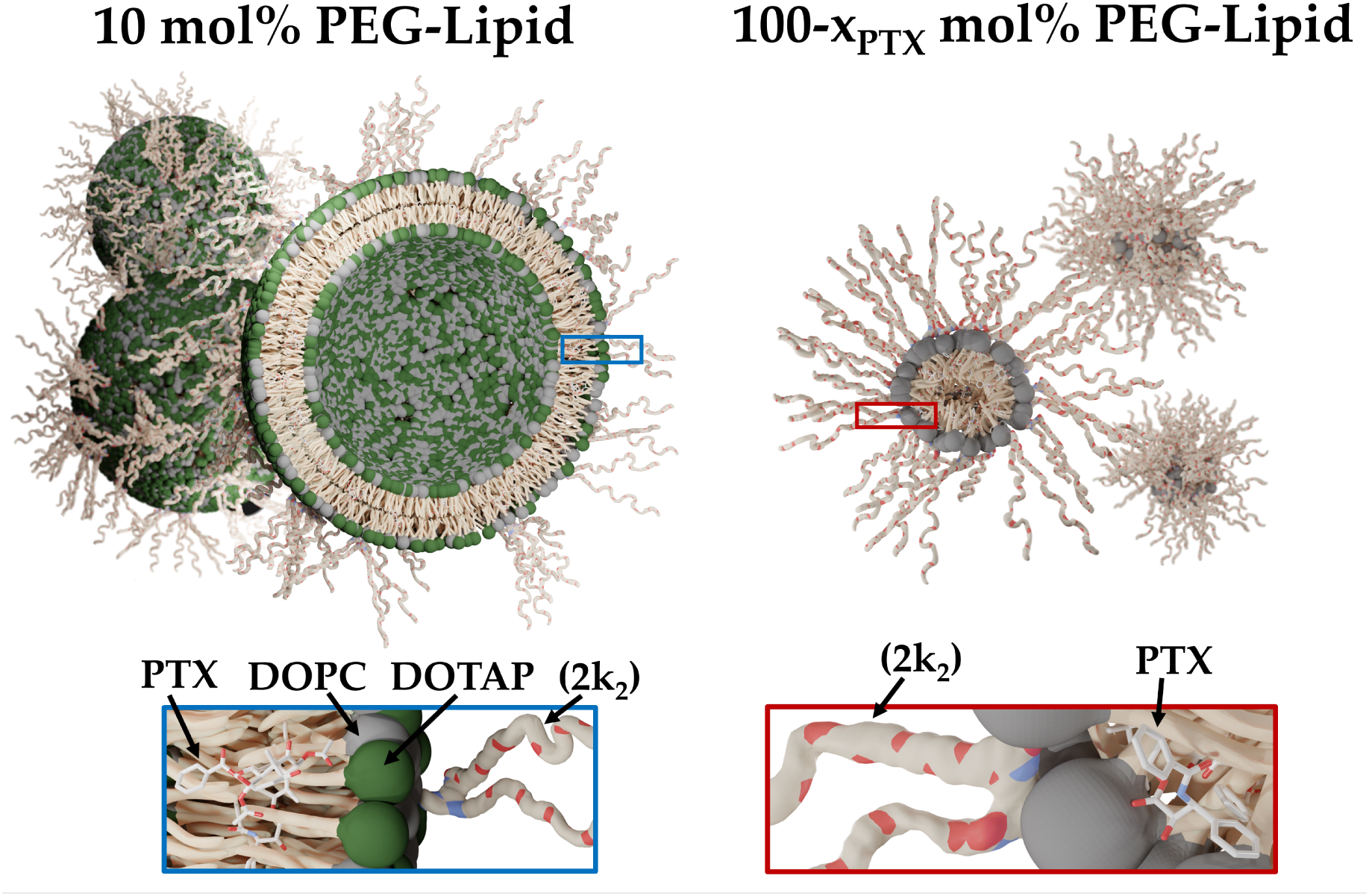
Graphical rendering of the PTX formulations formed using the custom high-curvature PEG-lipids synthesized in this study. The first formulation class examined (left half) results primarily in liposomes and consists of PEG-lipid, DOTAP, DOPC, and PTX in a 10:50:40–xptx:xptx mol ratio. The second formulation class examined (right half) results primarily in micelles and consists of PEG-lipid and PTX in a 100–xptx:xptx mol ratio. The PEG-lipid utilized in the formulation renderings is 2k_2_ (2 PEG2k chains). Owing to its hydrophobicity, PTX is solubilized entirely within the lipid membrane of the liposomal bilayer or micellar core. All lipids utilized in this study, including the custom-synthesized high-curvature PEG-lipids, contained monounsaturated dioleyl tails. Therefore, all lipid assemblies were expected to have membranes in the fluid (melted-chain) state.

## Methods

### Materials

ε-*N*-Boc-L-lysine methyl ester (Lys(BOC)-OMe) was purchased from Matrix Scientific. *N*- Boc-L-glutamic acid (Boc-Glu-OH) was obtained from TCI. N,N-Diisopropylethylamine was purchased from Chem Impex. Trifluoroacetic acid (TFA) and potassium hydroxide (KOH) was obtained from MilliporeSigma. 2-(1H-Benzotriazole-1-yl)-1,1,3,3-tetramethylaminium tetrafluoroborate (TBTU) was purchased from Oakwood Chemical. Paclitaxel (PTX) was obtained from Acros Organics as a powder. CellTiter 96 Aqueous-One Solution Cell Proliferation Assay was purchased from Omega. All solvents, acids, bases, and salts were obtained from Fisher Scientific. 1,2-dioleoyl-*sn*-glycerophosphatidylcholine (DOPC), 1,2-dioleoyl-3- trimethylammonium-propane (DOTAP) and 1,2-dioleoyl-*sn*-glycero-3-phosphoethanolamine-N- [methoxy(polyethylene glycol)-5000] (DOPE-5k) were purchased from Avanti Polar Lipids as powders. Dulbecco’s modified Eagle’s medium (DMEM) and penicillin/streptomycin were purchased from Invitrogen. Fetal bovine serum (FBS) was purchased from Gibco.

Stock solutions of PTX and lipids (both purchased and synthesized) were prepared in chloroform:methanol=3:1 (v/v) at concentrations of 1 mM (PTX, synthesized PEG-lipids, and DOPE-5k) or 10 mM (DOTAP, DOPC).

### Preparation of Fluid Lipid Nanocarriers

Lipid films were prepared by combining stock lipid components and PTX at the desired molar composition in a glass vial. Lipid mixtures were vortexed and the organic solvent evaporated to visual dryness using a stream of nitrogen. Lipid films were subsequently dried for 12 h in a vacuum pump. The resulting films were hydrated with high-resistivity water (18.2 MΩcm) to a working concentration of 1 mM total lipid (including PTX) and thoroughly vortexed.

### Kinetic Phase Diagrams (KPDs)

Following hydration and vortexing of fluid lipid PTX nanocarrier samples (designated t=0 h), the presence of PTX crystallization was evaluated with DIC imaging using a Nikon Eclipse Ti2 microscope equipped with 20x, 40x, and 60x objective lenses in addition to a 1.5x zoom lens. At set time points, 2 uL aliquots were taken from sample vials, placed on glass slides with a coverslip, and immediately imaged. New aliquots were taken for each time point. The first time point where imaging was performed was t=2 h following hydration, with imaging continuing at t=4, 6, 8, 10, 12, 24, 36, 48, 60, and 72 h time points. Sample vials were stored at room temperature during the experiment. Imaging was done at each time point in triplicates at minimum.

### Cell Culture

The human prostate cancer cell line PC3 (ATCC number: CRL-1435) was a gift from the Ruoslahti Lab (Burnham Institute, La Jolla). Cells were cultured in DMEM supplemented with 10% FBS and 1% penicillin/streptomycin. Cells were stored in an incubator at 37 °C infused with 5% CO_2_ in a humidified atmosphere. Passaging of cells was routinely performed upon reaching 80% confluency.

### Cell Viability Experiments

Samples in the 10 mol% PEG-lipid formulation class, as well as the unPEGylated No PEG samples, were processed post-hydration using tip sonication (Vibra-Cell^TM^, Sonics and Materials Inc., 30 W for 7 min). Samples in the 100–xptx mol% PEG-lipid formulation class were used post-hydration without further processing.

Cells were seeded onto 96-well plates at a density of 5,000 cells/well and final well volume of 100 uL supplemented DMEM. After 12 h, the media was replaced with media containing the PTX nanoparticle samples diluted at the designated final PTX concentration in DMEM. Following a 24 h incubation, the media was replaced with supplemented DMEM and the cells were incubated for a further 48 h. Following this, the media was replaced with CellTiter 96 AQueous- One Solution Cell Proliferation Assay reagent prepared as a 6x dilution in DMEM, with a final well volume of 120 uL. After incubating for 1 h, the absorbance at 490 nm was measured using a plate reader (Tecan Infinite 200 Pro). Each cell viability data point represents the average of four identically treated wells whose value was normalized to the viability of untreated cells.

### Statistical Analysis

All statistical analysis was performed in GraphPad Prism 8. Statistical significance between cell viability samples was calculated using Welch’s t-test. IC50 curves were generated by fitting to a four-parameter logistic equation.

## Results

### Synthesis of Customizable High-Curvature PEG-Lipids from Simple Building Blocks

The synthesis of the high-curvature PEG-lipids described in this study makes use of common building blocks which were conjugated in a combinatorial fashion to efficiently vary three key structural parameters (branching, charge, and PEG MW) (Scheme 1). The double-PEG-lipid compounds synthesized in this study differ significantly from commercially available PEG- phospholipids in the structure (and charge) of the lipid spacer. Therefore, we also prepared the neutral corresponding single-chain mPEG lipid (**2k**) and its cationic variant (**2k+**) as controls and to isolate any PEG branching-specific effects. Cationic functionalization (**1k_2_+, 2k+, 2k_2_+**) was performed by means of a lysine linker between the lipid tails and PEG chain(s). Oleyl tails (18:1) were used to ensure PTX lipid nanocarrier membranes were in the chain-melted (fluid) state at physiological temperature.^61^

**Scheme 1.**
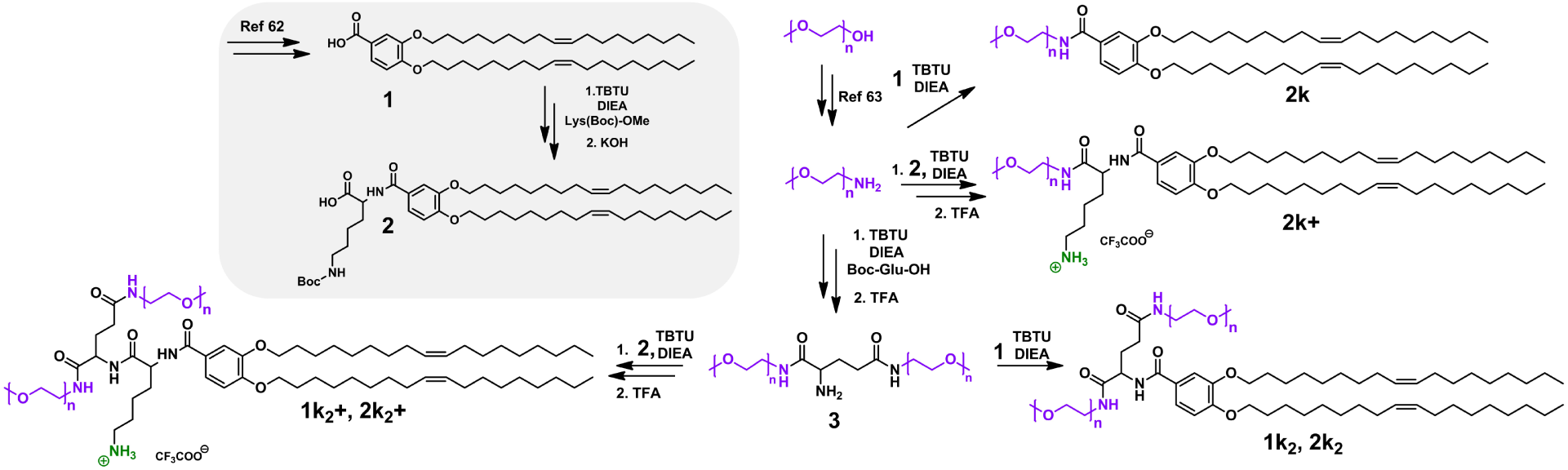
Synthesis of neutral and cationic high-curvature PEG-lipids with linear and double-PEG head groups. Boc-Glu-OH: *N*-Boc-l-glutamic acid; DIEA: N,N-diisopropylethylamine; Lys(Boc)-Ome: ε-*N*-Boc-l-lysine methyl ester; TBTU: 2-(1*H*-Benzotriazole-1-yl)-1,1,3,3-tetramethylaminium tetrafluoroborate.

The lipid tail building block **1** was synthesized as previously reported;^62^ from it, the cationic lipid building block **2** was synthesized by coupling with ε-*N*-Boc-L-lysine methyl ester followed by deprotection with KOH (Scheme 1, grey inset). Synthesis of the linear PEG building block was performed by conversion of mPEG-OH (MW of 1,000 g/mol [n≈22] or 2,000 g/mol [n≈44]) to mPEG-NH_2_ following a published protocol.^63^ The branched PEG building block **3** was afforded by coupling of mPEG-NH_2_ with *N*-Boc-L-glutamic acid, followed by Boc deprotection with TFA. The neutral single- (**2k**) and double- (**1k_2_, 1k_2_**) PEG-lipids were prepared by coupling of **1** to mPEG-NH_2_ or **3**, respectively. Similarly, the cationic linear (**2k+**) and branched (**1k_2_+. 2k_2_+**) PEG-lipids were prepared by coupling of **2** to mPEG-NH_2_ or **3**, respectively, followed by TFA deprotection.

### PTX Nanoparticle Formulations made from High-Curvature PEG-Lipids have Dramatically Improved PTX Solubility over Standard unPEGylated Formulations

Using DIC microscopy to observe formation of PTX crystals over time, we generated kinetic phase diagrams (KPDs) for the high-curvature PEG-lipids to evaluate their effect on PTX loading capacity across two formulation classes (Figures 3 and 4). The formulations in Figure 3 contain PEG-lipids at 10 mol%, together with DOTAP and DOPC. We employed mol ratios of DOTAP and DOPC similar to that in EndoTAG-1 (DOTAP:DOPC = 50:50–x_PTX_). We chose 10 mol% PEG-lipid as that ratio has been previously demonstrated to generate liposomes with a population of micelles.^15^ Additionally, 10 mol% PEG-lipid ensures the PEG chains are in the “brush” state on the surface of the membrane, which has been shown to lead to improved particle “stealth” properties *in vivo* compared to formulations with PEG-lipid in the mushroom state.^64, 65^ The formulations in Figure 4 contain PEG-lipids at 100–xptx mol%. We chose this ratio of PEG-lipid to promote predominant formation of micelles as previous studies using high-curvature lipids at similar ratios have demonstrated.^19, 40, 52, 54, 66^ For the formulations with 10 mol% PEG-lipid, we examined PTX solubility at 3, 5, 7, and 10 mol%. For the formulations with 100–xptx mol% PEG-lipid, we examined PTX solubility at 7, 10, and 13 mol% PTX.

**Figure 3.**
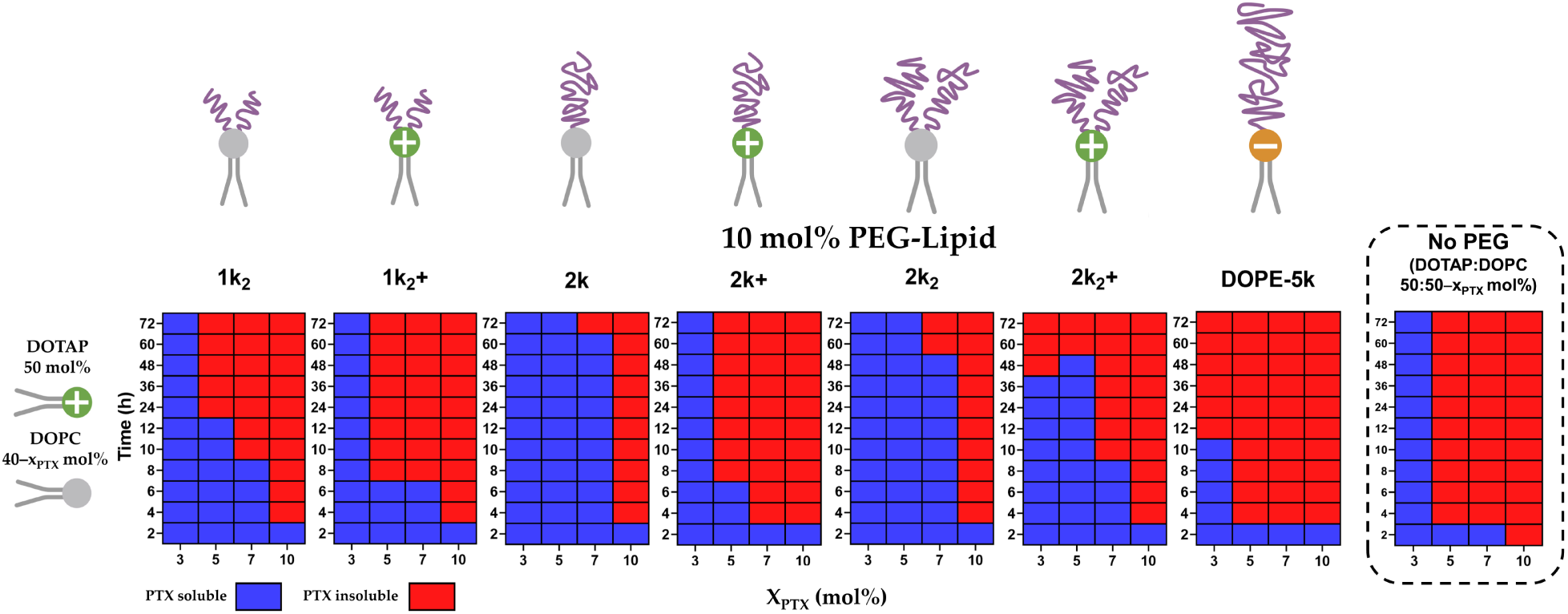
KPDs depicting PTX solubility over time for fluid lipid nanoparticle formulations containing high-curvature lipids at 10 mol%. The synthesized library of lipids allowed examination of the effect of charge (neutral or cationic), PEG length (1k to 5k), and PEG branching on PTX solubility. The commercially available DOPE-5k PEG-lipid and No PEG formulations were included as PEGylated and unPEGylated controls, respectively. PTX crystallization was assessed using DIC microscopy at the indicated time points, and blue and red color indicates the absence and presence of PTX crystals, respectively. Formulation composition: PEG-lipid:DOTAP:DOPC:PTX=10:50:40–xptx:xptx mol ratio or DOTAP:DOPC:PTX=50:50–xptx (No PEG). Time indicated refers to hours following hydration.

**Figure 4.**
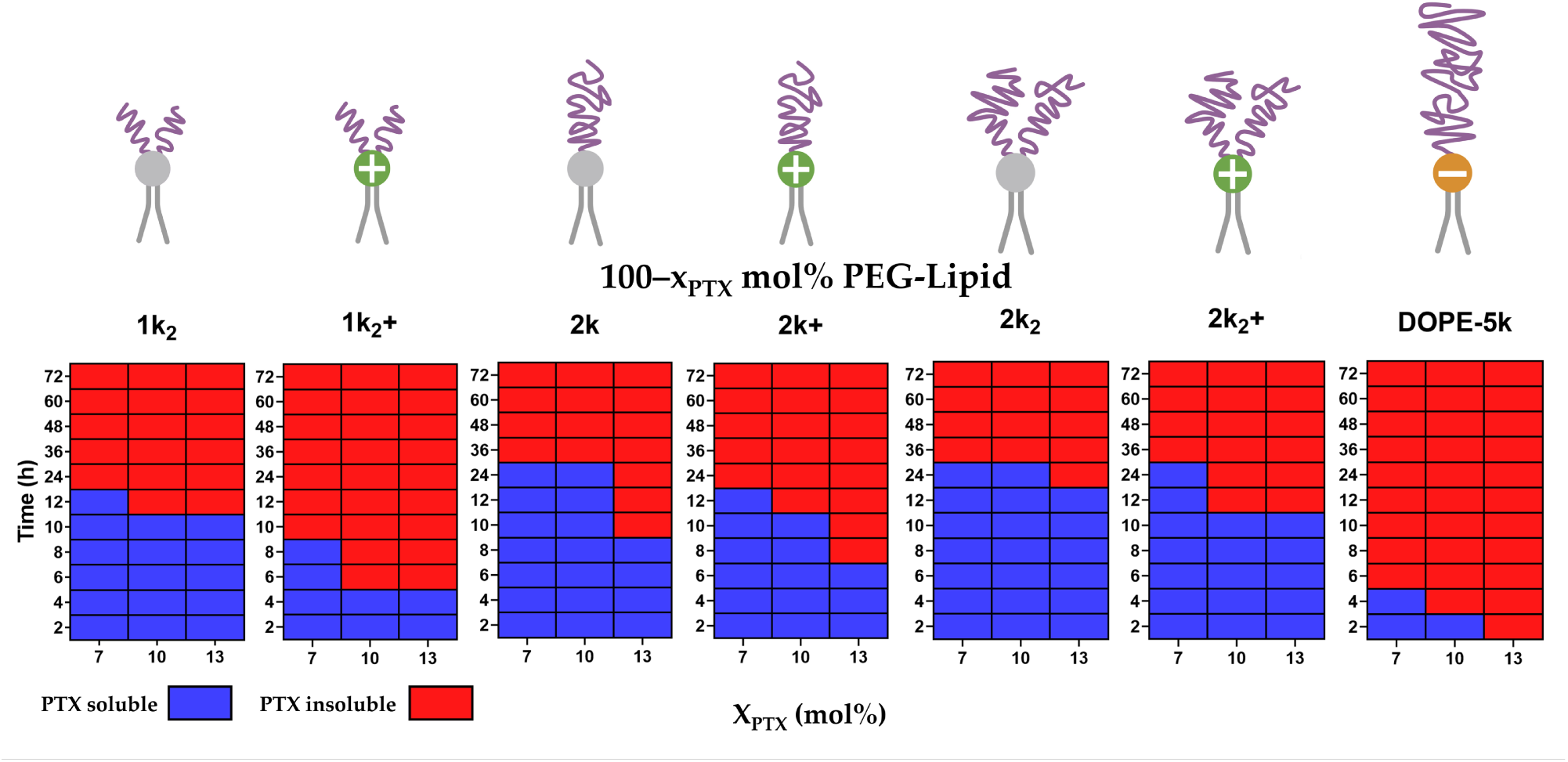
KPDs depicting PTX solubility over time for nanoparticle formulations containing high-curvature lipids at 100–xptx mol%. The commercially available DOPE-5k PEG-lipid was included as a control. PTX crystallization was assessed using DIC microscopy at the indicated time points, and blue and red color indicates the absence and presence of PTX crystals, respectively. Formulation composition: PEG-lipid:PTX=100–xptx:xptx mol ratio. Time indicated refers to hours following hydration.

The lowest PTX content evaluated within Figure 3, 3 mol%, represents the highest PTX loading ratio typically employed in unPEGylated fluid-membrane lipid nanoparticle PTX delivery systems, notably EndoTAG-1, which is currently undergoing clinical trials.^23^ At this PTX content, all the synthesized high-curvature lipids demonstrated solubility beyond 72 h, with the exception of 2k_2_+ and the commercially available DOPE-5k, which solubilized PTX for 48 and 12 h, respectively (Figure 3, two right-most diagrams). At 5 mol% PTX, only 2k and 2k_2_ maintained PTX solubility beyond 72 h. The PEG-lipids with a cationic charge exhibit poorer solubility of PTX compared to their neutral analogues (i.e. 1k_2_ and 1k_2_+, 2k and 2k+, 2k_2_ and 2k_2_+). At 7 mol% PTX, solubility further decreased for 1k_2_, 2k+, and 2k_2_+ while it stayed the same for 1k_2_+. Notably, while solubility also decreased for 2k and 2k_2_, these formulations were still able to stably solubilize PTX for a period of 72 and 60 h, respectively. The effect of branching on PTX solubility can be examined at 7 mol% PTX through comparison of the formulations containing 1k_2_ and 2k. 1k_2_ demonstrated significantly poorer PTX solubility compared to 2k despite a nearly identical total molecular weight of the PEG head group. At 10 mol% PTX none of the formulations maintained PTX solubility past 2 h. The unPEGylated No PEG formulation (which has an identical composition to EndoTAG-1 at 3 mol% PTX) displayed poorer PTX solubility over all PEG-lipid formulations with the exception of DOPE-5k, which had the poorest PTX solubility.

We chose to examine PTX kinetic solubility beginning from 7 mol% for the PEG-lipid:PTX=100–xptx:xptx formulations (Figure 4) due to our previous observations of high PTX loading in micelles formed by high-curvature lipids (unpublished data). At this ratio all samples remained stable for over 12 h, with the exception of the formulations prepared from 1k2+ and DOPE-5k. These samples showed crystals 10 h and 6 h after hydration, respectively. At 10 mol% PTX, the solubility of PTX decreased in all samples with the exception of formulations prepared from 2k and 2k_2_, which remained stable up to 36 h. The change in PTX membrane solubility between 7 and 10 mol% PTX is more drastic for the cationic PEG-lipids compared to their neutral analogues: crystals appeared 12 h faster at 10 mol% PTX for 2k+, and 24 h faster for 1k_2_+ and 2k_2_+, compared to no change for 2k, 12 h faster for 1k_2_, and no change for 2k_2_. At 13 mol% PTX, PTX solubility further decreased for all samples except for the formulation prepared from 1k_2_. At this high loading, PTX crystallized from the DOPE-5k-containing formulations immediately after hydration. Remarkably, the formulations prepared from 1k_2_, 2k_2_, and 2k_2_+ demonstrate prolonged PTX solubility even at 13 mol%. The lipid 2k_2_ particularly solubilizes PTX well, not forming PTX crystals until 24 h after hydration (compared to 12 h for 1k_2_ and 2k_2_+), long enough for practical clinical applications. For the PEG-lipid=100–_PTX_ formulations, PEG branching results in marginally improved PTX solubility at 13 mol%, showing PTX solubility until 12h for 1k_2_ compared to 10 h for 2k. Inclusion of the cationic spacer disrupts PTX membrane solubility more than the improvement provided by branching, as 1k_2_+ formulations form PTX crystals at 6 h compared to 8 h for the formulations containing the straight chain 2k+ with an analogous total molecular weight of the PEG head group.

Notably, all newly synthesized lipids demonstrate strikingly higher PTX solubility in both formulation classes compared to the commercially available DOPE-5k (Figures 3 and 4, rightmost panels). These findings are contrary to previous reports of PEG-lipid hindering PTX loading and clarify that the effect of PEG-lipids on PTX solubility can be positive or negative depending on the PEG-lipid’s structure.^52^ The absolute extent of PTX solubility was found to be higher in the formulations containing PEG-lipid at a high ratio (100–xptx mol%, Figure 4) compared to those at a low ratio of 10 mol% (Figure 3). Consistent to both formulation classes, samples containing PEG-lipids with 1kDa PEG chains exhibited lower PTX solubility than those containing PEG-lipids with 2kDa PEG chains. The addition of a cationic charge in the spacer hinders PTX solubility in nearly all cases. Branching of the PEG head groups (i.e. double-PEG head groups) somewhat decreases PTX solubility in most cases where a direct comparison of lipids with identical total chain length (1k_2_ and 2k, 1k_2_+ and 2k+) is possible, with the exception of 100–xptx 1k_2_+ formulations (better solubility at 7–13 mol% PTX over 2k+ formulations) and 10 mol% 1k_2_+ formulations (better solubility at 7 mol% PTX over 2k+ formulations). We found the greatest PTX solubility among the 10 mol% PEG-lipid formulations in samples containing 2k and 2k_2_ at 7 mol% PTX (72 and 60 h, respectively, Figure 3). Within the 100–xptx mol% formulations we found the greatest PTX solubility in samples containing 2k_2_ at 13 mol% PTX (24 h, Figure 4). Within the context of PEG chain branching on PTX solubility, the excellent PTX solubility of the double-PEG-lipid 2k_2_ formulations are noteworthy.

### High-Curvature PEG-Lipids show Potent Cytotoxicity against Human Cancer Cells when used in Micelle-Inducing Formulations

We next examined the cytotoxicity of formulations containing the newly synthesized high-curvature PEG-lipids at 10 mol% (Figure 5) and 100–xptx mol% (Figure 6a), shown as green bars. In both figures, we included a sample with PTX in the absence of a carrier (PTX Only, white bars) as a control. In Figure 5 we additionally included an unPEGylated formulation (No PEG, gray bars) to represent the formulation for EndoTAG-1 (DOTAP:DOPC:PTX=50:47:3 mol%). The human pancreatic cell line PC3 was used to screen the high-curvature PEG-lipids, as its cell viability has been previously shown to be highly responsive to varying PEG-lipid mol%.^15^ The cytotoxicity experiments depicted in Figure 5 and Figure 6 were performed at a final well concentration of 20 nM PTX. We chose this concentration as it is close to the IC50 of a 2kDa PEG-lipid formulation previously reported.^15^ In order to assess the effect on cytotoxicity owing to differences in PTX solubility, we chose to perform cytotoxicity experiments at the low and high extremes of the PTX mol% ratios explored in the KPDs (Figures 3 and 4). For the 10 mol% PEG-lipid formulations, we evaluated cytotoxicity at 3 mol% and 10 mol% PTX (Figure 5, left and right graphs, respectively). For the 100–xptx mol% PEG-lipid formulations, the PTX ratios were 7 mol% and 13 mol% (Figure 6a, left and right graphs, respectively). We chose to exclude DOPE-5k formulations owing to their poor PTX solubility as shown in Figures 3 and 4. In a subsequent experiment, we examined the cytotoxicity of formulations containing 2k_2_ and DOPE-5k (PEG-lipid:PTX=93:7 mol%) at varying final PTX concentrations (Figure 6b). The PEG-lipid:PTX=93:7 mol% formulation for 2k_2_ was chosen as it exhibited the most potent cytotoxicity at 20 nM PTX (Figure 6a, left graph).

**Figure 5.**
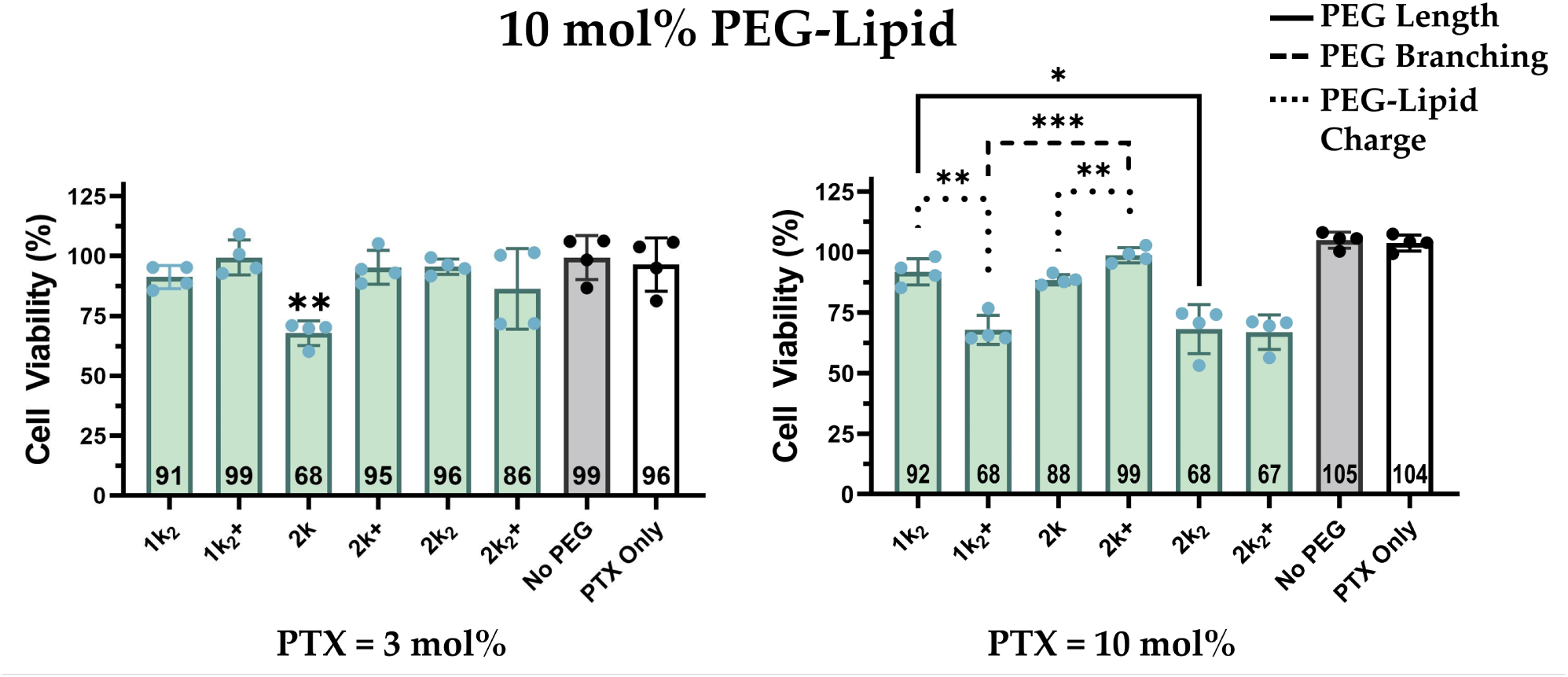
Cytotoxicity of PTX-loaded formulations containing high-curvature PEG-lipids at 10 mol%. Cytotoxicity was evaluated at 3 mol% PTX (left graph) and 10 mol% PTX (right graph) loading ratios against PC3 cells. In addition to the high-curvature PEG-lipid formulations (green bars), an unPEGylated formulation (No PEG, gray bars) and free PTX (PTX Only, white bars) were included as controls. Formulations were added to cells at a final [PTX]=20 nM. Formulation composition: PEG-lipid:DOTAP:DOPC:PTX=10:50:40–xptx:xptx mol ratio; DOTAP:DOPC:PTX=50:50–xptx:xptx mol ratio (No PEG sample). Lipid films were hydrated and immediately processed with tip sonication, followed by addition to cells. Cell viability was normalized to untreated cells. Cell viabilities shown represent the mean ± SD (n=4). Statistical significance was calculated using Welch’s t-test, where *p<0.05, **p<0.01, ***p<0.001. The p value for the 2k sample in the 3 mol% PTX group (left graph) was calculated compared to the No PEG and PTX Only samples. Bracket lines between samples (right graph) tested for statistical significance were labeled as follows: solid line for samples differing in PEG length, dashed line for samples differing in PEG branching, dotted line for samples differing in PEG-lipid charge.

**Figure 6.**
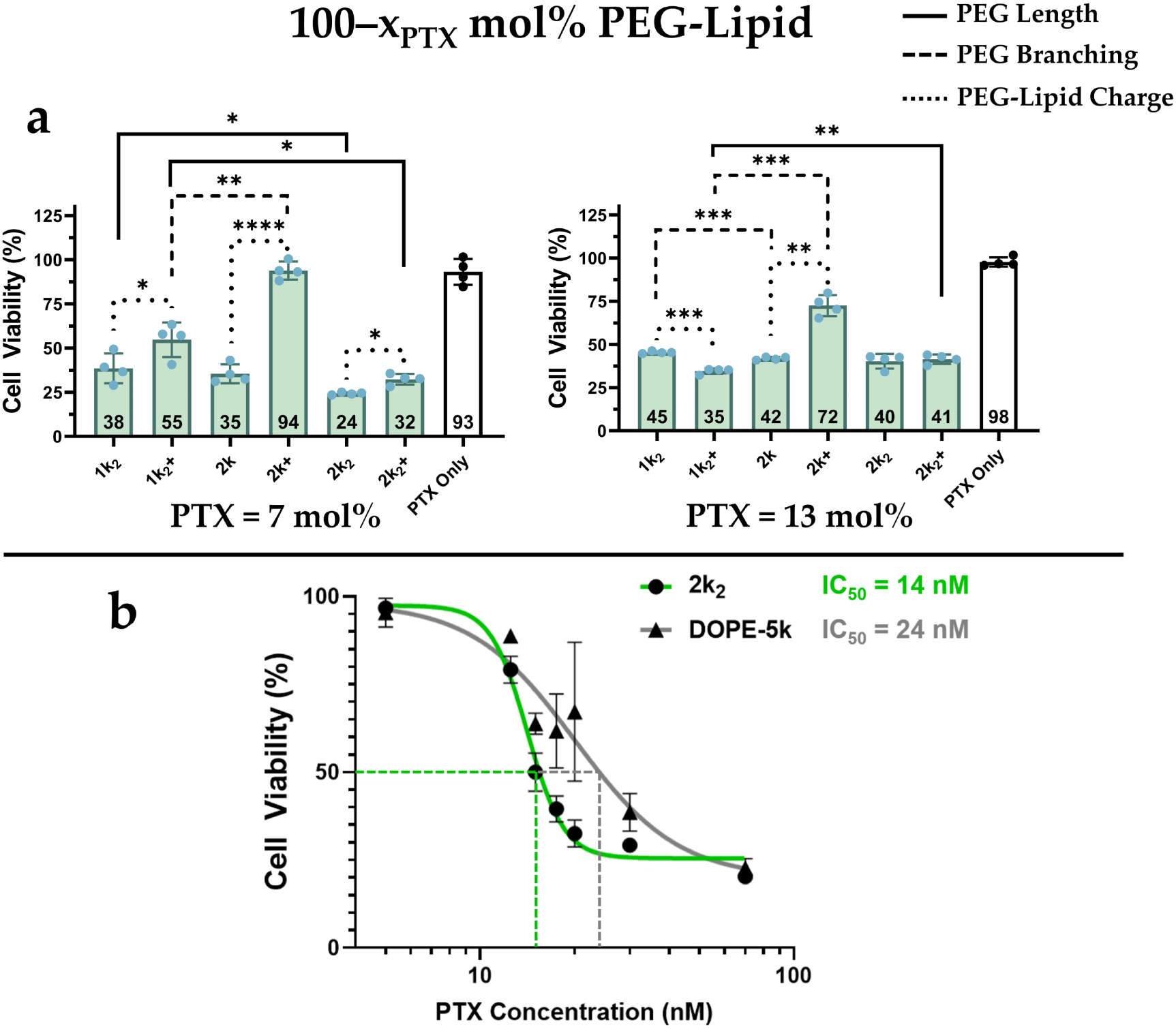
Cytotoxicity of PTX-loaded formulations containing high-curvature PEG-lipids at 100–xptx mol%. (a) Cytotoxicity was evaluated at 7 mol% PTX (left graph) and 13 mol% PTX (right graph) loading ratios against PC3 cells. In addition to the high-curvature PEG-lipid formulations (green bars), PTX without a lipid carrier (PTX Only, white bars) was included as a control. Formulations were added to cells at a final [PTX]=20 nM. Statistical significance was calculated using Welch’s t-test, where *p<0.05, **p<0.01, ***p<0.001, ****p<0.0001. Bracket lines between samples tested for statistical significance were labeled as follows: solid line for samples differing in PEG length, dashed line for samples differing in PEG branching, dotted line for samples differing in PEG-lipid charge. (b) IC50 curves generated for formulations containing 2k_2_ (green curve) and DOPE-5k (gray curve). Formulation composition: PEG-lipid:PTX=93:7 mol ratio. Cell viability is normalized to untreated cells. Cell viabilities shown represent the mean ± SD (n=4).

Within the 10 mol% PEG-lipid formulations (Figure 5) at a PTX loading ratio commonly employed in PTX lipid nanoparticles (3 mol%, left graph),^21, 67, 68^ we found that none of the high-curvature PEG-lipids showed significantly higher cytotoxicity compared to the PTX only (white bar) and unPEGylated (gray bar) samples, with the exception of the 2k formulation (p<0.01, cell viability = 68%). Greater differences between the samples emerge in the 10 mol% PTX loading group (Figure 5, right graph). We found the 1k_2_+, 2k, 2k_2_, and 2k_2_+ samples to have significantly higher cytotoxicity (68%, 88%, 68%, and 67%, respectively) compared to the No PEG (105%) and PTX Only (104%) samples (Dunnet’s multiple comparisons test, not shown on graph). Within the 10 mol% PTX loading group, an increase in PEG chain length from 1k_2_ to 2k_2_ led to improved cytotoxicity (92% to 68%, p<0.05), while the effect from an increase in length from 1k_2_+ and 2k_2_+ was not found to be significant (68% and 67%). We determined PEG branching to not significantly affect cytotoxicity between the 1k_2_ and 2k formulations, respectively. However, we found cytotoxicity in the double-PEG-lipid 1k_2_+ formulation to be lower than the single-PEG-lipid 2k+ formulation (68% and 99%, p<0.001). Addition of a cation to the lipid head group led to opposing effects on cytotoxicity; 1k_2_+ had improved cytotoxicity compared to 1k_2_ (68% and 92%, p<0.01), while 2k+ had reduced cytotoxicity compared to 2k (99% and 88%, p<0.01). Furthermore, we found 2k_2_ and 2k_2_+ to display similar cytotoxicities (68% and 67%).

Comparing between samples within the 100–xptx mol% PEG-lipid formulation class with a 7 mol% PTX (Figure 6a, left graph), an increase in PEG chain length from 1k_2_ to 2k_2_ resulted in enhanced cytotoxicity (38% to 24%, p<0.05), as did the increase in length from 1k_2_+ and 2k_2_+ (55% and 32%, p<0.05). We found PEG branching to not significantly affect cytotoxicity between 1k_2_ and 2k samples, although cytotoxicity was higher in the double-PEG-lipid 1k_2_+ compared to the single-PEG-lipid 2k+ (55% and 942%, p<0.01). Cationic PEG-lipids in this group all demonstrated poorer cytotoxicity compared to their neutral PEG-lipid analogues: 38% and 55% for 1k_2_ and 1k_2_+ (p<0.05), 35% and 94% for 2k and 2k+ (p<0.0001), and 24% and 32% for 2k_2_ and 2k_2_+ (p<0.05). The 13 mol% PTX group within the 100–xptx mol% PEG-lipid formulation class represented the highest PTX loading ratio examined in this study (Figure 6a, right graph). The samples within this group all had statistically more potent cytotoxicity than the PTX Only control (Dunnet’s multiple comparisons test, not shown on graph). We found an increase in PEG chain length from 1k_2_+ to 2k_2_+ to slightly reduce cytotoxicity (35% and 41%, p<0.01), with no significant difference in the cytotoxicities of the neutral analogues 1k_2_ and 2k_2_. PEG branching led to slightly poorer cytotoxicity between 1k_2_ and 2k (45% and 42%, p<0.001). However, cytotoxicity was higher in the branched 1k_2_+ compared to the straight chain 2k+ (35% and 72%, p<0.001). We determined PEG-lipid cationic charge to have opposing effects on cytotoxicity depending on the sample, with an improvement to cytotoxicity from 1k_2_ to 1k_2_+ (45% and 35%, p<0.001), a reduction in cytotoxicity from 2k to 2k+ (42% and 72%, p<0.01), and nearly no difference between 2k_2_ and 2k_2_+.

Taken as a whole, within the 100–xptx mol% PEG-lipid formulation class, formulations in the 13 mol% PTX group displayed poorer cytotoxicity compared to the 7 mol% PTX group. This finding is in agreement with the PTX solubility elucidated in the KPDs (Figure 4), where we found nearly all samples loaded with 7 mol% PTX (with the exception of DOPE-5k which was excluded from cytotoxicity experiments) to stably encapsulate PTX on the order of 24 hr. In contrast, in the 13 mol% PTX formulation group only 2k_2_ could stably solubilize PTX for 24 hr. The greatest potency among the samples was displayed by 2k_2_ at 7 mol% PTX, supporting the argument that stable PTX solubilization within lipid carriers is critical to achieving maximal drug delivery. Among the 7 mol% and 13 mol% PTX loading groups, cytotoxicity consistently increased with longer PEG lengths. The effects of PEG branching between these two PTX loading ratios were similar: both 7 mol% and 13 mol% PTX-containing samples showed nearly no change in cytotoxicity between 1k_2_ and 2k formulations, while formulations containing the double-PEG-lipid 1k_2_+ had dramatically higher cytotoxicity than those with the single-PEG-lipid 2k+. Formulations made from 2k+ had markedly poorer cytotoxicity than all other PEG-lipid samples in the 100–xptx mol% PEG-lipid group. The reason for the poor cytotoxicity of samples using 2k+ in this micellar formulation is unclear, although it’s relatively poor PTX solubility as revealed by the KPDs was likely a contributing factor. A clear trend of hindered cytotoxicity among cationic PEG-lipids compared to neutral PEG-lipids is present in the low PTX (7 mol%) loading formulation group (Figure 6a, left graph). This trend is less defined in the 13 mol% PTX group (Figure 6a, right graph). Thus, it is evident that although PTX solubility is important in predicting cell cytotoxicity performance, other factors act in conjunction to influence cytotoxicity, such as the enhancement in cellular uptake and cytotoxicity of cationic particles compared to neutral ones.^69^

In a final experiment, we determined the IC50 of 2k_2_ in its most potent formulation (100– x_PTX_ mol% PEG-lipid, x=7 mol%) and compared it to the IC50 of the commercially available DOPE- 5k (Figure 6b). We found a sharper dose-response curve for 2k_2_ compared to DOPE-5k in addition to a lower IC50 (14 nM and 24 nM PTX, respectively). This result confirms the high PTX loading and delivery efficiency of fluid micellar nanoparticle formulations, particularly those made with the high-curvature double-PEG-lipid 2k_2_ custom-synthesized in this study.

## Discussion and Conclusion

Despite the improved tumor penetration and blood stability of lipid micelles compared to liposomes, lipid micelle PTX delivery systems employing PEG remain limited in efficacy by their poor PTX solubility.^40, 52, 53^ Prior studies have attained high PTX loading in fluid lipid nanocarriers by strategies such as adding lysolipids to diacyllipid formulations or mixing unsaturated and saturated lipids to form nanostructured lipid carriers, achieving up to 12–15 mol% PTX loading.^40, 70^ However, these delivery methods suffer from drawbacks such as the instability of lysolipids in blood. Additionally, such high PTX loading strategies have exclusively utilized liposomes rather than micelles and often display suboptimal performance despite encapsulating high ratios of PTX, with one study finding a higher IC50 and comparable *in vivo* pharmacokinetics to Taxol.^70^

The KPD and cytotoxicity data generated in this study for formulations employing the 6 custom-synthesized high-curvature PEG-lipids provide insight into the relationship between PTX solubility and effective PTX delivery across two main parameters, 1) fluid lipid nanoparticle structure (liposomal or micellar), and 2) PEG chain length, PEG-lipid charge, and PEG branching (single- or double-PEG head groups). The two formulation classes explored in this study, 10 mol% PEG-lipid and 100–xptx mol% PEG-lipid formulations, generate populations of fluid lipid nanocarriers that are primarily liposomes and primarily micelles, respectively.^15, 40^ Comparison of PTX solubility between these classes reveals a drastic higher PTX solubility in PEGylated fluid lipid micelles compared to liposomes. While all samples within the 10 mol% formulation group showed PTX crystallization within 4 h when encapsulating 10 mol% PTX (Figure 3), the group consisting primarily of micelles (100–xptx mol% PEG-lipid formulations) could stably solubilize PTX at 10 mol% for at least 12 h in nearly all cases (Figure 4). We found micellar formulations made from the neutral double-PEG-lipid 2k_2_ to result in the greatest PTX solubility in this study, encapsulating 13 mol% PTX for 24 h when used at 100–xptx mol%. An interesting finding is that in addition to the consistent improvement in PTX loading among samples when going from a low to high PEG-lipid ratio, the relative ability of different samples to solubilize PTX was maintained (i.e. samples showing highest PTX solubility at 10 mol% PEG-lipid also showed the highest PTX solubility at 100–xptx mol% PEG-lipid). This result supports the argument that the high-curvature PEG-lipids synthesized in this study, and not DOTAP or DOPC, were the primary sources for the improvement in PTX solubility over unPEGylated samples such as EndoTAG-1.

By examining the KPDs and cell viability data for all the synthesized high-curvature PEG-lipids, we were able to establish trends regarding the effect of PEG chain length, PEG-lipid charge, and PEG branching (single or double PEG chains) on PTX kinetic solubility and downstream cytotoxicity. Longer PEG chain lengths displayed improved PTX solubility and cytotoxicity in nearly all cases. In contrast, addition of a positive charge to the synthesized PEG-lipids resulted in lower PTX solubility over neutral PEG-lipid counterparts in all cases. The reduced PTX solubility of cationic PEG-lipids corresponded to poorer cytotoxicity in many of the formulations (with other cases having nearly equal cytotoxicities). The influence of PEG branching was more elusive; among some formulations we found double-PEG-lipids to result in more potent cytotoxicity over single-PEG-lipids with roughly identical molecular weights, where within other formulations, double-PEG-lipids displayed nearly-identical cytotoxicities.

Interestingly, within the 10 mol% PEG-lipid formulation class, many samples showed higher cytotoxicity at the high PTX loading ratio (10 mol%) over the low PTX loading ratio (3 mol%, Figure 5) having an identical final well [PTX] = 20 nM. This is a surprising finding as no formulation loaded with 10 mol% PTX maintained PTX solubility past 4 hr (Figure 3). Additionally, PTX solubility in these samples were further reduced due to all the formulations in Figure 5 being sized-down using tip sonication prior to their use in cytotoxicity experiments, as tip sonication has been observed to reduce PTX solubility for DOTAP/DOPC/2kDa PEG-lipid formulations.^15^ Therefore, it is highly likely the samples loaded with 10 mol% PTX all contained PTX crystals before being added to cells. One possible mechanism for the unexpected cytotoxicity enhancement in the 10 mol% PTX-loaded formulations is the relative enrichment in high-curvature PEG-lipids over 3 mol% PTX-loaded samples, as formulations containing 10 mol% PTX had a 7 mol% lower ratio of DOPC. Given DOPC has C_0_ ≈ 0, a higher relative ratio of PEG-lipid: DOTAP and DOPC within the 10 mol% PTX group may have resulted in a larger population of micelles than in the 3 mol% PTX group, improving overall cytotoxicity despite onset of PTX crystallization.

Another curious finding was that the commercially available PEG-lipid DOPE-5k demonstrated starkly lower PTX solubility in both formulation classes compared to the 6 PEG-lipids synthesized in this study. A previous study investigating micelle PTX carriers made from DOPE-5k reported a solubility limit in agreement with our KPDs (Figure 4).^40^ However, given our observed trend of improved PTX solubility with longer PEG chain lengths, the poor solubility of formulations containing DOPE-5k is unexpected. Two structural differences between DOPE- 5k and the 6 PEG-lipids synthesized in this study provide possible rationales for this observation: namely that DOPE-5k is anionic (where the 6 synthesized PEG-lipids are neutral or cationic), and that DOPE-5k contains a glycerol linker and phosphate moiety in the head group (where the synthesized PEG-lipids have a linker based on 3,4-dihydroxybenzoic acid). To our knowledge no anionic lipid carrier PTX formulations other than DOPE-5k micelles have been previously investigated. However, anionic liposomal formulations of another extremely hydrophobic drug, all-trans-retinoic acid, have achieved a high drug encapsulation efficiency (99%) and multi-month membrane drug stability.^71^ Additionally, a study examining liposomal delivery of the hydrophobic drug curcumin found significantly higher drug encapsulation in anionic liposomal formulations compared to cationic formulations (87.8% and 57.5%, respectively).^72^ These results are in contrast to the KPDs of liposomal formulations in this study (Figure 3), which uncover notably poorer PTX solubility in formulations made from the anionic DOPE-5k compared to formulations made from the cationic 1k_2_+, 2k+, and 2k_2_+. Therefore, while it is clear the solubility of hydrophobic drugs are impacted by membrane charge, whether drug solubility is enhanced or diminished in cationic or anionic lipid carriers varies depending on the drug. Similarly, modulation of lipid linker structure (including length, bulk, and charge) can have substantial effects on membrane properties.^73, 74^ This is illustrated by one study that demonstrated a change in gene delivery efficiency from being potently efficient to nearly transfection incompetent when the orientation of the ester linkages in the cationic lipid tails was reversed.^75^ Examining the structure of the 6 synthesized PEG-lipids, we expect the aromatic ring linking the two oleyl tails and PEGylated head group to lie near the membrane-water interface when self-assembled into particles. The presence of the rigid aromatic ring at the interface is expected to lead to persistent “gaps” in the membrane interface, particularly in the 100–xptx mol% PEG-lipid formulations where the PEG-lipid is the sole lipid component. These “gaps” may allow for stable solubilization of PTX molecules, allowing favorable nonpolar interactions between the PEG-lipid’s aromatic ring and the hydrophobic regions of PTX, while simultaneously positioning PTX molecules close enough to the membrane interface to permit hydrogen bonding between the hydroxyl, ester, and amide groups of PTX and water.

Given we found PEG chain branching to have a minimal impact on PTX solubility and cytotoxicity, it is intriguing that formulations made from the double-PEG-lipid 2k_2_ demonstrated the highest absolute PTX solubility and cytotoxicity. Employing the trends uncovered in this study allows us to formulate a rationale for this discovery. First, the relatively long PEG chain lengths of 2k_2_ (in comparison to 1k_2_ or 1k_2_+) translates to improved PTX solubility and cytotoxicity. Second, the absence of charge on 2k_2_ further improves PTX solubility and cytotoxicity compared to the cationic PEG-lipids. Finally, branching may have effects on particle stability that were not directly measured within this study but nevertheless led to potent cytotoxicities. An example of such an effect would be higher cellular uptake of formulations employing double-PEG-lipids (i.e. 2k_2_) compared to straight chain PEG-lipids (i.e. 2k) as a result of enhanced serum stability of branched PEG-lipids. A recent study exploring the pharmacokinetics of doxorubicin-loaded liposomes incorporating double-PEG-lipids lends support to this possibility, with the authors finding a significantly lower immunogenic response and higher *in vivo* tumor efficacy resulting from double-PEG-lipid formulations compared to straight chain PEG-lipids.^76^

This study elucidated the tremendous therapeutic potential of fluid lipid micellar nanoparticles formulated from a library of custom-synthesized high-curvature PEG-lipids. We expect this work to broadly invigorate further exploration of micellar nanoparticle carriers utilizing high-curvature PEG-lipids for not only the delivery of PTX, but other hydrophobic drugs as well. One compelling progression is the modification of the PEG-lipids with functional ligands such as tumor-homing peptides on the PEG extremities, adding a layer of active targeting functionality to the micellar nanoparticles. Here we find the branched PEG-lipid 2k_2_ particularly of value as it can be conjugated to two identical or unique ligands, locally tethering them in space.

## Acknowledgements

This research was supported by the National Institutes of Health under award R01GM130769 (mechanistic studies on developing lipid nanoparticles for drug delivery). Partial support was provided by the National Science Foundation under award DMR-1807327 (membrane phase behavior of lipid nanoparticles).

## Notes

### Competing Interest Statement

The authors have declared no competing interest.

